# Evolution of Replication Origins in Vertebrate Genomes: Rapid Turnover Despite Selective Constraints

**DOI:** 10.1101/411470

**Authors:** Florian Massipa, Marc Laurent, Caroline Brossas, José Miguel Fernández-Justel, María Gómez, Marie-Noelle Prioleau, Laurent Duret, Franck Picardb

## Abstract

**Background:** The replication programme of vertebrate genomes is driven by the chromosomal distribution and timing of activation of tens of thousands of replication origins. Genome-wide studies have shown the frequent association of origins with promoters and CpG islands, and their enrichment in G-quadruplex sequence motifs (G4). However, the genetic determinants driving their activity remain poorly understood. To gain insight on the functional constraints operating on replication origins and their spatial distribution, we conducted the first evolutionary comparison of genome-wide origins maps across vertebrates.

*Results:* We generated a high resolution genome-wide map of chicken replication origins (the first of a bird genome), and performed an extensive comparison with human and mouse maps. The analysis of intra-species polymorphism revealed a strong depletion of genetic diversity on an ~ 40 bp region centred on the replication initiation loci. Surprisingly, this depletion in genetic diversity was *not* linked to the presence of G4 motifs, nor to the association with promoters or CpG islands. In contrast, we also showed that origins experienced a rapid turnover during vertebrates evolution, since pairwise comparisons of origin maps revealed that only 4 to 24% of them were conserved between any two species.

**Conclusions:** This study unravels the existence of a novel genetic determinant of replication origins, the precise functional role of which remains to be determined. Despite the importance of replication initiation activity for the fitness of organisms, the distribution of replication origins along vertebrate chromosomes is highly flexible.

## INTRODUCTION

During each cell cycle, the genome must be accurately replicated to ensure the faithful transmission of the genetic material to daughter cells. In vertebrates, as in many other eukaryotes, this fundamental process is initiated at specific sites, called replication origins (Prioleau and MacAlpine 2016). Several studies in humans and mice have shown that their genomes consist of large domains (from 200 kb to 2 Mb) that are replicated at different time points of S-phase (Farkash-Amar et al. 2008, Hansen et al. 2010, Hiratani et al. 2008). This spatio-temporal programme of genome replication is driven by the chromosomal distribution of replication origins and by their timing of activation. Thanks to the development of new techniques, based on the purification of short nascent DNA sequences (SNS), it has been possible to obtain genome-wide maps of replication origins at the kb resolution in human (Besnard et al. 2012, Cadoret et al. 2008, Picard et al. 2014), mouse (Almeida et al. 2018, Cayrou et al. 2015), Drosophila (Comoglio et al. 2015) nematode (Pourkarimi et al. 2016, Rodriguez-Martinez et al. 2017), and the human parasite *Leishmania major* (Lombrana et al. 2016). In humans and mice, 65,000 to 250,000 origins have been identified (Almeida et al. 2018, Besnard et al. 2012, Cayrou et al. 2015, Picard et al. 2014), which reflects the number of origins that are active in a population of cells. However, autoradiographic evaluation of the distance between two active origins (Berezney et al. 2000), has led to the estimate that, at most, 30, 000 origins are actually activated in each cell cycle in mammals. Furthermore, the comparison of 5 different human cell types has uncovered substantial plasticity in the spatial programme of replication: 10~ 15% of replication origins are cell-type-specific, whereas 35 ~60% are constitutively active (Besnard et al. 2012, Picard et al. 2014).

Despite these important advances in the characterization of replication origins, the genetic determinants of their activity are still poorly understood. In both human and mice, replication origins are strongly enriched in transcription start sites (TSSs) and CpG islands (CGIs) (Cadoret et al. 2008, Cayrou et al. 2015, Picard et al. 2014, Sequeira-Mendes et al. 2009). They are also associated with specific epigenetic marks (Julienne et al. 2015, Picard et al. 2014), depending on the progression of the cell cycle. Mammalian replication origins are enriched in G-rich motifs that are capable of forming DNA secondary structures called G-quadruplexes (G4) (Besnard et al. 2012, Cayrou et al. 2012, Maizels and Gray 2013, Picard et al. 2014) and there is experimental evidence that G4 elements can directly contribute to the function of origins (Valton et al. 2014). However, the presence of a G4 motif is not sufficient for origin activity (Valton et al. 2014), and 50% of these motifs are found outside of known origins (Maizels and Gray 2013), implying that other genetic elements must contribute to the origin activity.

To gain insight into the selective constraints that operate at the location of replication origins and the genetic elements that determine their activity, we decided to investigate the evolution of replication origins in vertebrates. For this purpose, we mapped origins in the chicken genome. We chose chicken because the evolutionary distance between mammals and birds is such that only functionally constrained elements are conserved between their genomes (Wallis et al. 2004). Here, we report a comparative analysis of replication origin landscapes across mammals and birds, and an investigation of selective constraints that act on their sequences, based both on an inter-species comparison and on an analysis of intra-species genetic polymorphism.

The polymorphism analysis demonstrated that the locus of maximal SNS enrichment within replication origins contained functionally important sequence elements that were clearly subject to purifying selection and distinct from G4 motifs. However, inter-species comparisons revealed a poor conservation of these elements on a larger evolutionary scale and showed that the location of replication origins evolved rapidly. Furthermore, the chromosomal distribution of origins in mice is very different from that observed in human and chicken. These observations indicate that despite the importance of replication initiation activity for the fitness of organisms, the distribution of replication origins along chromosomes is highly flexible.

## RESULTS

### Large-scale variations in replication initiation landscapes

To study the evolution of replication initiation landscapes, we generated the first map of replication origins in a bird genome, by sequencing short nascent strands (SNS) in the chicken DT40 cell line (derived from a bursal lymphoma, see Methods). For comparison with mammalian origins, we reanalysed all previously published human and mouse SNS datasets (Almeida et al. 2018, Besnard et al. 2012, Cayrou et al. 2015, Picard et al. 2014) using the same detection pipeline based on scanning windows and family-wise error rate control, with a resolution of 500 bp (Picard et al. 2014). This method detects genomic segments on which the accumulation of reads is significantly higher than the background, which depends on the global read depth of the experiment (Table S1). Given that the sensitivity of origin detection depends on the sequencing depth (Fig. S1), we sub-sampled SNS reads so that all three species reached the same sequencing depth (see Methods section). In human and mouse, SNS datasets are available for different cell lines (human: IMR90, H9, K562, HeLa and iPS; mouse: mESC10, MEFs). To limit differences due to tissue-specific origin activity, human/mouse comparisons are based on origins detected in H9 and mESC10 cell lines, both of which correspond to embryonic stem cells.

As expected given the differences in genome sizes (1.2 Gb in chicken vs.~ 3 Gb in mammals), we identified fewer origins in chicken (33, 500) than in human (155, 395) or mouse (205, 881) (Table S1). However, variations in the number of origins were not entirely explained by genomes size since we observed a 2.5-fold variation in origins density, from 34 origins per Mb in chicken to 84 origins per Mb in mouse (55 origins per Mb in human). Mouse origins are on average shorter (690 bp), than human (791 bp) or chicken origins (891 bp), but the short size of mouse origins might be due to the selection of shorter nascent strand in mouse experiments (0.5 to 2 kb) compared to chicken (1.5 to 2.5 kb) and human (1 to 2 kb) experiments. Overall, replication origins cover 3% (chicken), 5.5% (human), and 4.3% (mouse) of the genome.

Visual comparison of several homologous regions in the three species revealed that replication initiation landscapes were quite different in mouse compared to human and chicken: we observed many more peaks in mouse, but these peaks were less intense (Fig. 1). To extend this comparison over the whole genome, we analysed the distribution of replication initiation activity in 100 kb windows (Fig. 2). In mouse, 5% of the genome concentrates 26.7% of the replication initiation activity, compared to 49.7% in human and 65.3% in chicken. Thus, origin activity is more uniformly distributed in mouse than in human and chicken (Fig. 2). It should be noted that for both human and mouse, we obtained consistent results with SNS datasets corresponding to distinct cell lines generated by different laboratories (Fig. S2).Thus, the differences in replication initiation landscapes observed between human and mouse were reproducible, and were not explained by the cell type variability. Interestingly, the more uniform replication initiation landscape that we observed in mouse compared to human coincided with a more uniform GC content distribution (Waterston et al. 2002). In agreement with previous observations in human (Besnard et al. 2012, Cadoret et al. 2008), we observed strong positive correlations between the origin density and regional genomic GC content in all three species (Fig. S3-(A-C). Indeed, our observation of differences in replication landscapes between mouse and human or chicken vanished after correcting for GC content differences among species (Fig. S3-(D-E)). All these observations indicate that the number of replication origins is not simply dictated by the evolution of genome size and that even within mammals, there are substantial differences in the distribution of replication initiation activity along chromosomes.

**FIG. 1:**
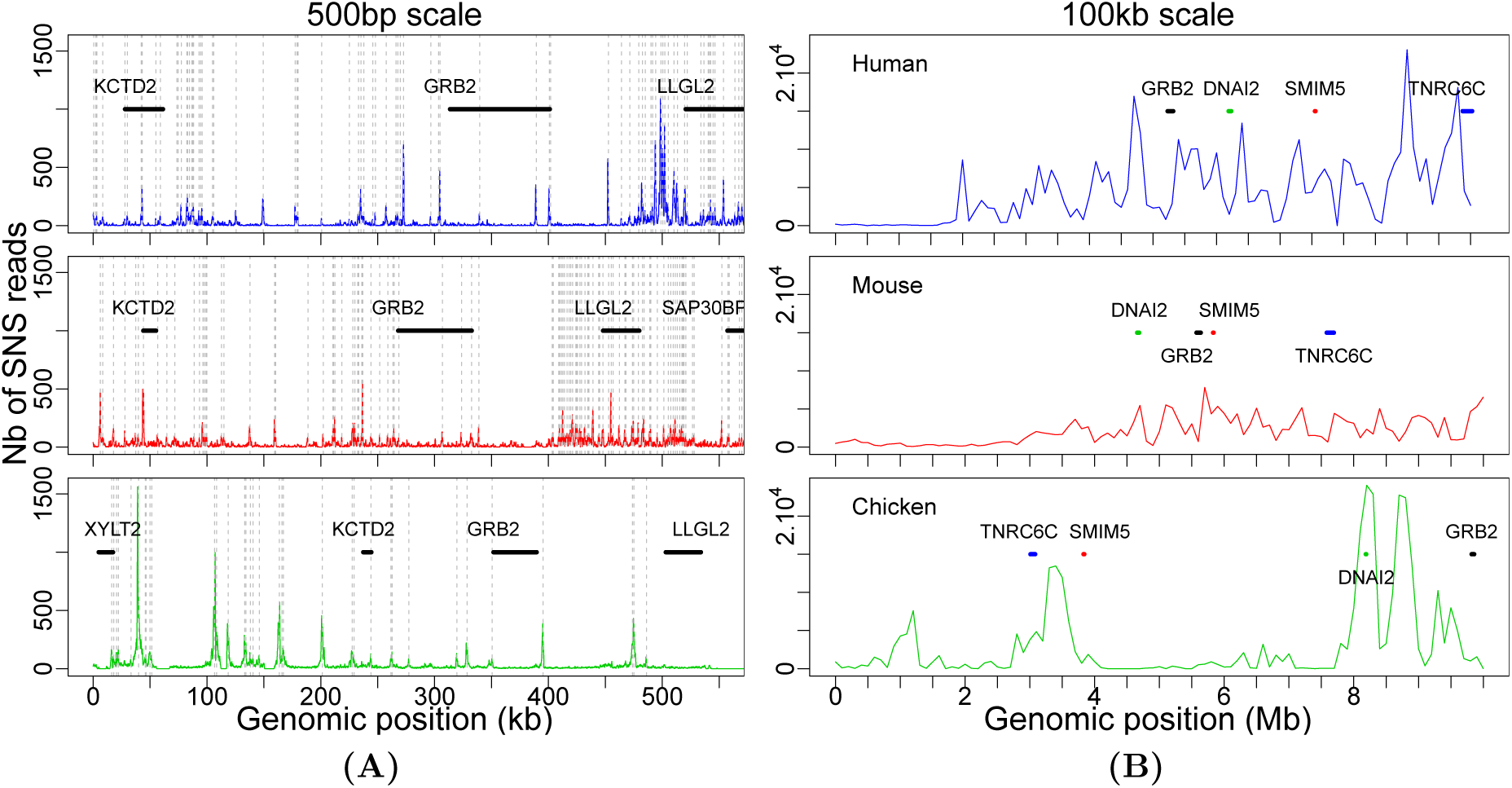
Profile of replication initiation activity along a representative homologous locus in human (top), mouse (middle) and chicken (bottom) genomes. **(A)** Number of SNS reads per 500 bp window in human chr17:73,000,600-73,550,600, mouse chr11:115,376,000-115,926,000, and chicken chr18:10,486,119-11,036,119 regions. **(B)** Number of SNS reads in 100 kb windows in a 10 Mb region centred on the region presented in **(A)**. Vertical grey dotted lines represent detected origins, and several homologous genes are annotated with thick lines.

**FIG. 2:**
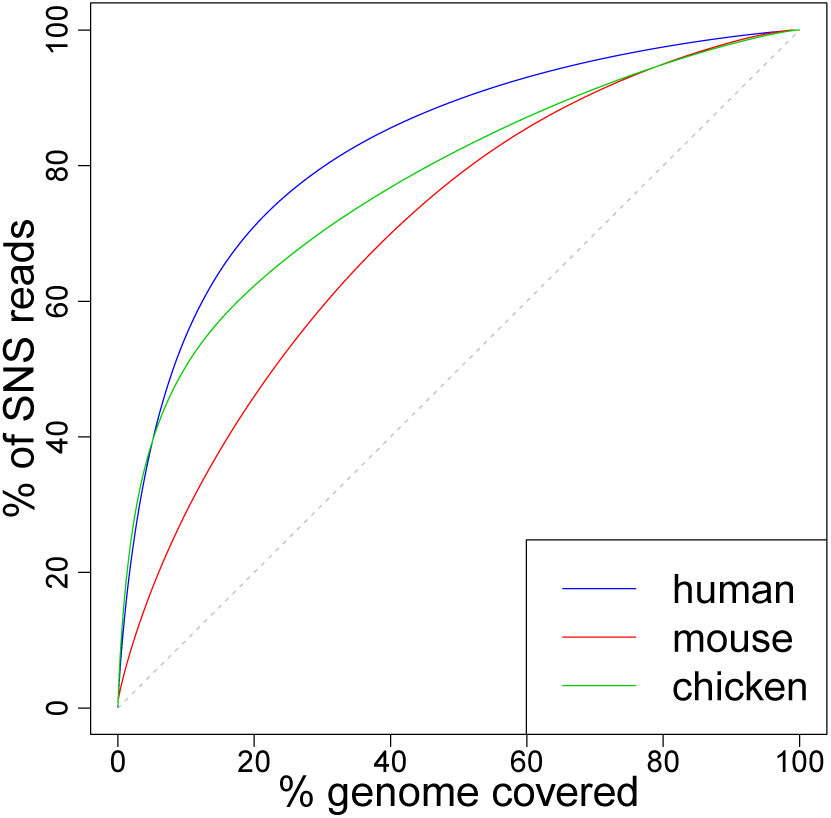
Replication initiation landscapes in vertebrates: Cumulative proportion of SNS reads mapped to the *x* richest 100 kb regions of the genomes. The black broken line corresponds to the expectation under a uniform distribution of reads.

### High precision mapping of replication origins

To gain insights into the genetic elements controlling replication initiation, we analysed the average patterns of sequence composition and selective constraints along origins. One key point in these analyses is to define a relevant reference point to align origins. Here we chose SNS peak, i.e., the point of maximal SNS enrichment along the sequence (see Methods section). It has been previously shown that replication origins are associated with specific base composition skew profiles (Cayrou et al. 2012, 2011), likely resulting from the inversion of mutation patterns at the transition between leading and lagging strands (Haradhvala et al. 2016, Touchon et al.2005). Interestingly, we observed that in the three species, both GC and AT-skew profiles (*S*_GC_ = (G *-* C)*/*(G + C), *S*_AT_ = (A *-* T)*/*(A + T)) invert exactly at the SNS peak position (Fig. 3). Our fine-scale analysis at kb resolution allowed us to map the position of maximal skew *~* 50 bp to the SNS peak (compared with 280 bp identified in mouse (Cayrou et al. 2012)), suggesting that our method identifies the position of the replication start site very precisely.

**FIG. 3:**
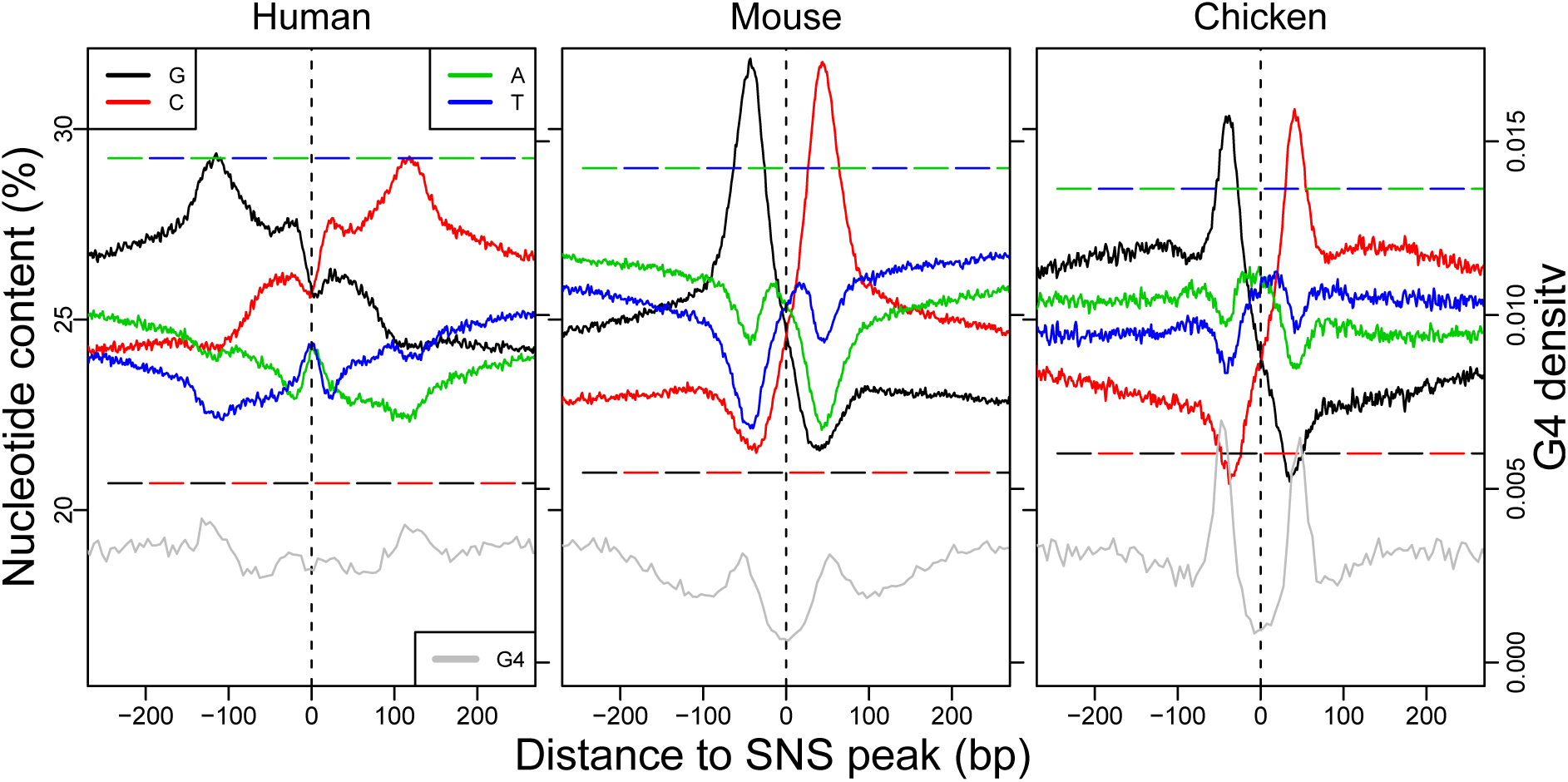
Replication origin sequence signature: Average nucleotide content in the 500 bp region centred on the SNS peak (plain line). Genome-wide average nucleotide contents are indicated by broken lines. Grey lines present the total number of G4s found in all origins for 5 bp windows. To achieve a bp resolution for G4s, we summarize each G4 by the position of its middle nucleotide.

### Polymorphism data show evidence of selective pressure on replication origins

To investigate selective constraints impacting replication origins, we first analysed intraspecific sequence polymorphism. In humans, we used data from the 1000 genomes project (based on 2500 individuals), focusing on single nucleotide polymorphisms (SNPs) for which the nature of the ancestral and derived alleles could be inferred by comparison with great apes genomes (1000 Genomes Project Consortium 2015). To detect signatures of selection, we analysed SNPs with different derived allele frequencies (DAFs). Common variants (DAF > 10%) generally correspond to relatively old mutations that have spread in the population as a result of genetic drift, selection or GC-biased gene conversion (gBGC) (Glemin et al. 2015). Rare variants (DAF < 1%) predominantly correspond to recent mutations, which are less affected by selection or gBGC and hence are expected to more accurately reflect the prevalence of mutational events (Kimura and Ohta 1973). To distinguish the effect of selection from that of gBGC, we separately analysed SNPs resulting from GC→ AT mutations, AT → GC mutations or GC-conservative mutations. For rare variants, we observed that the SNP density was constant around SNS peaks and corresponded to the genome-wide average (Fig. 4). This finding suggested that the mutation rate within origins did not differ from the rest of the genome. In contrast, we observed a strong deficit in common variants, specifically in the ~40 bp region surrounding the SNS peak. This deficit is observed for all categories of SNPs, both GC→ AT and AT→GC (Fig. 4-(C-D)), or GC-conservative (Fig. S4), which implies that it is not caused by gBGC. Similarly, we observed a deficit of common indels (Fig. S4-(C-D)) in the same 40 bp region. Hence, these observations demonstrated that the immediate vicinity of the SNS peak was subject to purifying selection. Many replication origins are associated with CGIs or TSSs, i.e., functional regulatory elements that are under selective pressure to control gene expression. Interestingly, the signature of selection around the SNS peak was observed for all categories of origins, irrespective of the presence of CGIs or TSSs (Fig. 4-(A-B)). Together with the observation that the SNP depletion occurred specifically in the immediate vicinity of the SNS peak, this result indicated that the signature of selection was directly linked to the function of replication initiation and not to the fact that origins are often associated with other functional elements.

**FIG. 4:**
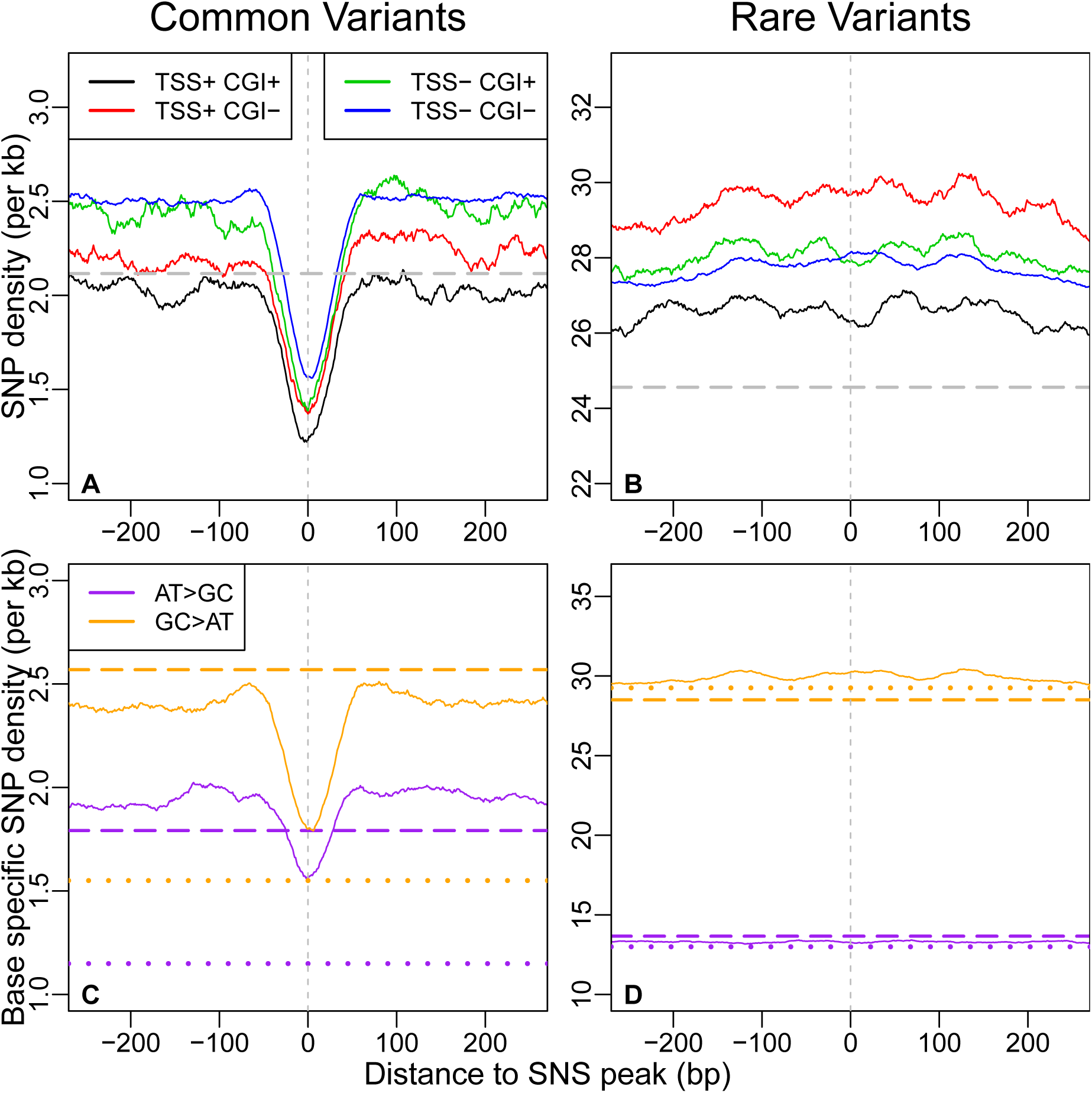
Density of genetic polymorphism around origins: average SNP density in the vicinity of all human SNS peaks (plain lines) for different classes of origins, depending on their association with CGI and/or TSSs. Genome-wide averages are indicated by broken lines. **(A)** Common variants (*DAF* > 10%). **(B)** Rare variants (*DAF* < 1%). **(C-D)** Base-specific SNP density (see Methods section) in the vicinity of all human SNS peaks (plain line). Broken lines: genome-wide average. Dotted line: average base-specific SNP density in all human coding exons. **(C)** Common A or T to C or G (AT > CG) and A or T to C or G (AT > CG) variants (*DAF* > 10%). **(D)** Rare variants (*DAF* < 1%).

**FIG. 5:**
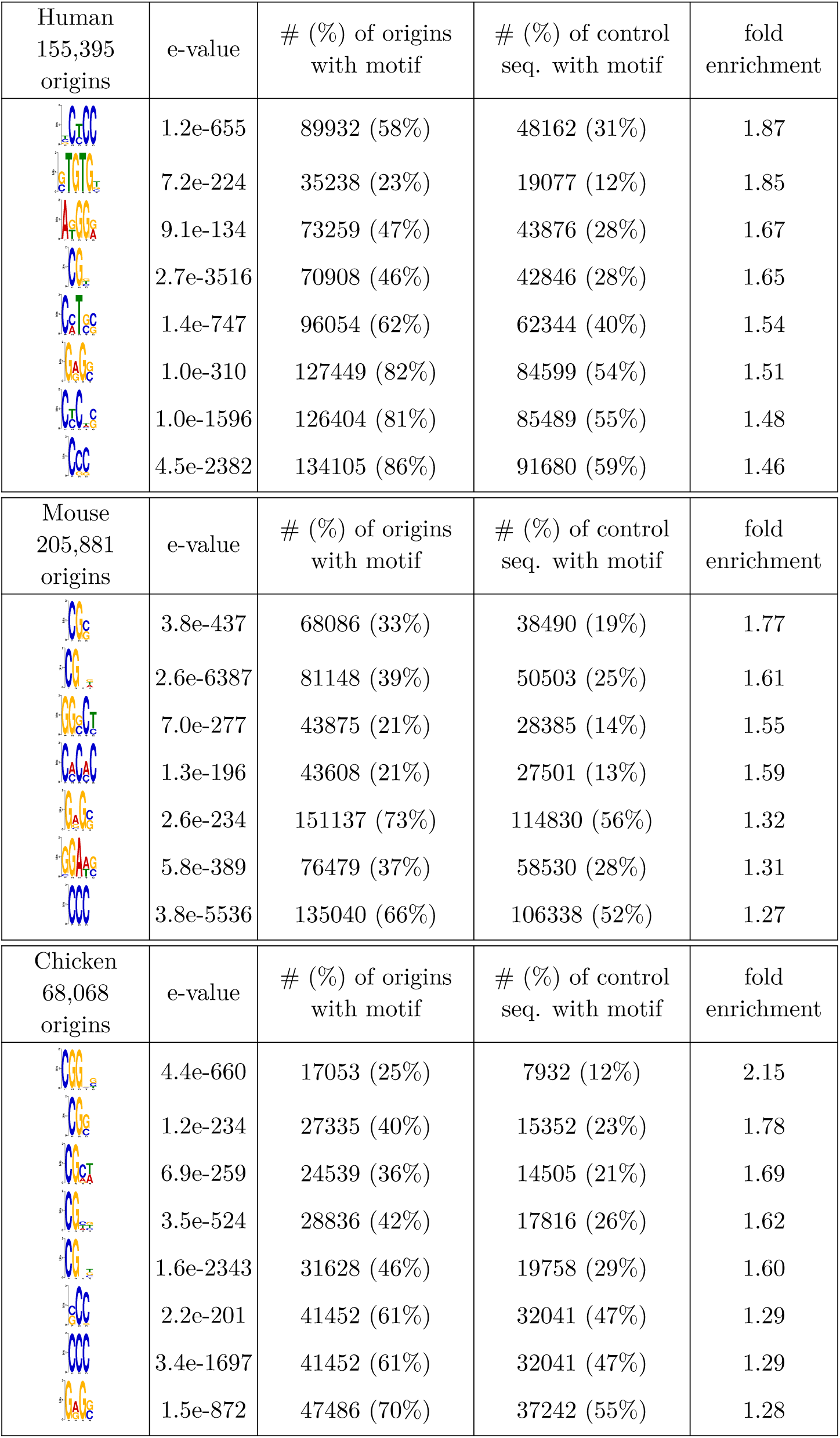
Motifs enriched in the 40 bp core region of origins for each species. Motifs were detected by adjusting the background model to compositional heterogeneities (see Methods). Control sequences were obtained by randomly sampling the same number of 40 bp regions in each genome.

Overall, the density in common SNPs within SNS peaks was 25% lower than in flanking regions (Fig. 4-A). Although this deficit was weaker than the 40% depletion observed in coding exons (Fig. 4-(C), dotted line), it clearly indicated that the sequence of human replication origins was subject to quite strong purifying selection. Hence, the core region of replication origins included genetic elements that are important for their function.

In mouse and chicken, polymorphism datasets are based on relatively small population samples, which does not allow the comparison of rare and common variants. However, profiles of SNPs density also revealed a specific signature in the~ 40-bp region centred on the SNS peak (Fig. S5), which suggested that origins are subject to similar selection patterns in the three species.

### Selective pressure on replication origins is driven by sequence features that differ from G4s

Previous analyses of human and mouse replication origins have revealed an enrichment in G4 sequence motifs (Besnard et al. 2012, Cayrou et al. 2012), and there is experimental evidence that these elements can contribute to their function (Valton et al. 2014). However, in all three species, we observed that the regions of the highest G4 density did not coincide with the central 40-bp region under selective pressure (Fig. 3). This result suggested that this selective pressure was not targeted towards G4s and that the 40-bp core region of origins probably contained another class of functional elements.

To search for new sequence signatures of the 40-bp core region of origins, we investigated motif enrichment using randomly selected 40-bp segments as a control. Since the compositional heterogeneities of genomes strongly impacted motif detection, the random segments were first clustered based on their kmer composition (5-mers) using *k*-means, hence providing clusters of homogeneous composition (see Methods, Supp. Table S2), in which the motif search could be performed. We next annotated each core region of origins to a background cluster depending on their 5-mer contents and then searched for motifs cluster-wise to adjust the procedure to a heterogeneous background. Using the DREME software in the MEME suite (Bailey 2011), we selected motifs with strong e-values (≤1e^-200^) and further inspected motifs that were most enriched in origins compared with random segments. The detected motifs were all very short (≤ 6 bp), and strong enrichments were detected cluster-wise in motifs that were rich in CG dinucleotides for all species (Table 5). The AC (GT) repeats previously identified in mouse (Cayrou et al. 2015) were confirmed and detected in human origins with 1.85-fold enrichment. Then, CCC(GGG) trinucleotides emerged as a shared sequence signature among species, with a two-fold cluster-wise enrichment in origins compared with random sequences. Since we showed an average depletion of G-quadruplexes within the constrained regions (Fig. 3), these trinucleotides were likely to be distinct from flanking G-quadruplexes. Finally, the GAG (CTC) motif was another common feature between the three species. Consequently, our motif analysis highlighted common short sequence features within this evolutionarily constrained 40-bp region, with functional roles that remain to be experimentally investigated.

### Replication origins have experienced a rapid turnover in vertebrates

Given the selective constraints that were identified in humans, we further analysed the conservation of replication origins on a larger evolutionary scale. We first investigated whether origins detected in chicken, human or mouse overlapped with conserved genomic segments (CGSs), identified in pairwise whole-genome alignments between these species (Fig. 6-(A-B)). To estimate the number of overlaps expected by chance, we randomly sampled a set of segments in the mappable regions of the human genome with the same size and chromosomal distribution as origins (see Methods section). We found that approximately 25% of human origins and 21% of mouse origins overlapped with chicken-mammal CGSs (Table 1). In both cases, these features corresponded to a 2-fold enrichment compared to random expectation.

**TABLE 1:** Conservation of origins across species. Top: Percentage of origins that overlap conserved genomic segments (CGS). Bottom: Percentage of functionally conserved origins. In parentheses: Conservation enrichment compared to the control experiment (see Randomization procedures in the Methods section for more details).

| % of origins overlapping a CGS |  |  |  |
| --- | --- | --- | --- |
|  | Human | Mouse | Chicken |
| Human |  | 72.6% (**×1.2**) | 24.8% (**×2.1**) |
| Mouse | 75.7% (**×1.2**) |  | 20.6% (**×2.1**) |
| Chicken | 48.9% (**×1.7**) | 40.7% (**×1.6**) |  |
| % of origins functionally conserved |  |  |  |
| Human |  | 20.7% (**×2.4**) | 7.9% (**×4.5**) |
| Mouse | 15.6% (**×2.5**) |  | 4.3% (**×4.0**) |
| Chicken | 23.7% (**×3.4**) | 18.2% (**×2.8**) |  |

**FIG. 6:**
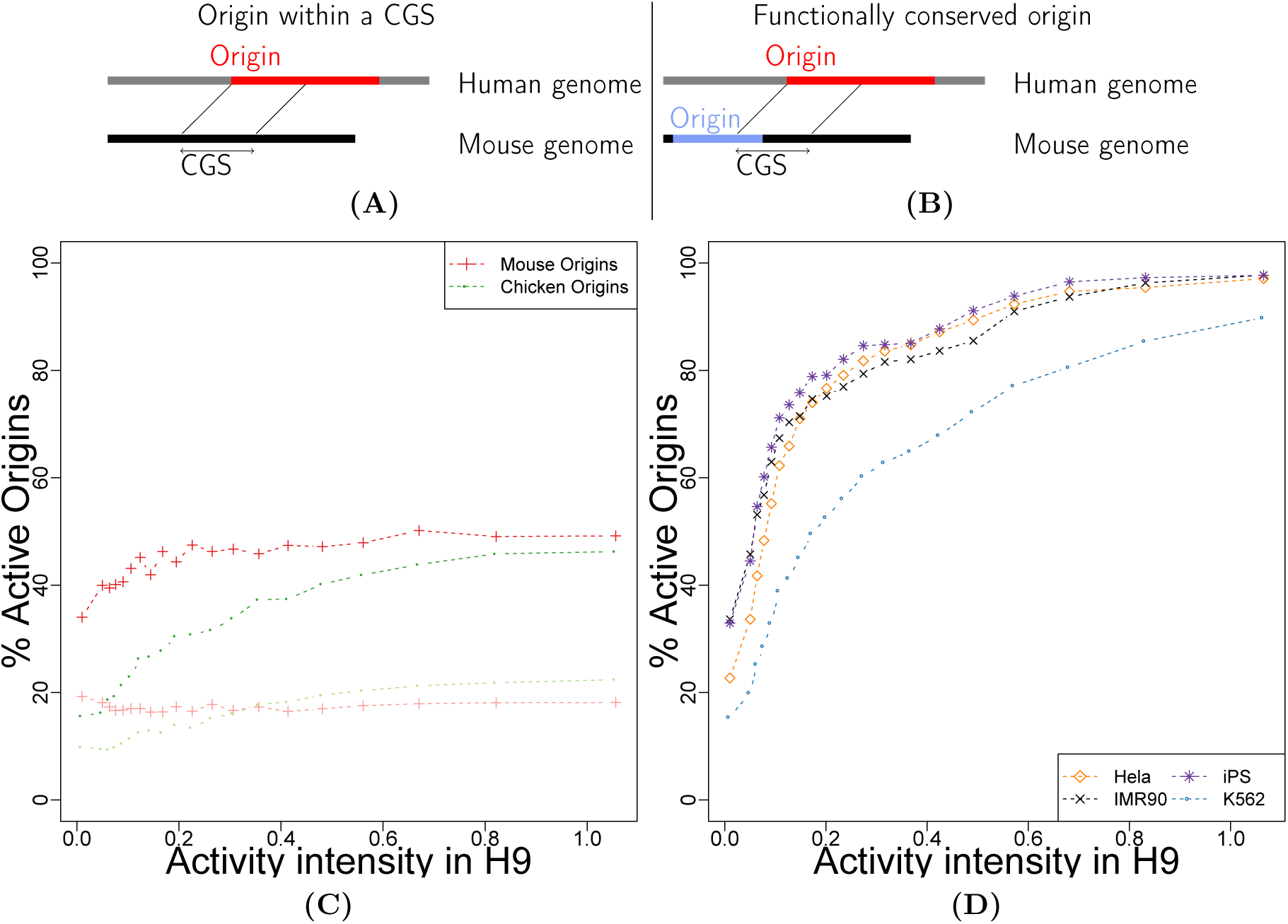
Analysis of functional conservation of human origins. Schematic representation of **(A)** origins within CGS and **(B)** functionally conserved origins. **(C)** Functional conservation of human origins among origins within mammal-bird CGSs: Percentage of origins conserved in chicken and mouse. Light colour: Random chicken and mouse genomic segments resembling origins (see the Methods section). **(D)** Percentage of origins active in different human cell types after coverage correction (see the Methods section).

We then assessed the functional conservation of origins and found that 8% of human origins were functionally conserved in chicken (1.8% expected). Similarly, 4.3% of mouse origins were functionally conserved in chicken (1.1% expected). Hence, only a small fraction of origins appeared to be functionally conserved across vertebrates.

Estimates of the level of functional conservation might be subject to several artefacts. First, we observed that the fraction of functionally conserved origins increased with their level of activity (Fig. 6-C and Fig. S6), which might reflect limitations in the sensitivity of detection of the less active origins. Second, the analysis of distinct cell types in chicken from those analysed in mammals might also contribute to an underestimation of the fraction of functionally conserved origins. To check these two hypotheses, we took advantage of data available in five different human cell lines, focusing on origins overlapping a CGS only. We observed that among the strongest human origins, more than 90% were detected in all cell types (Fig. 6-D), which implies that the detection sensitivity was very high for this subset of origins and *ii*) most of them were constitutive. Interestingly, among the strongest origins overlapping a chicken-mammal CGSs, only 45% were functionally conserved in chicken (Fig. 6-C). Hence, the relatively low level of functional conservation detected between human and chicken origins was not due to limited sensitivity, nor to cell-type specificity issues.

Overall, 8% of human origins have been conserved both in terms of sequence and function since the divergence of mammals and birds. For a comparison we analysed the conservation of TSSs of protein-coding genes: 50% of human TSSs overlapped with chicken-human CGSs, among which 43% corresponded to TSSs annotated in the chicken genome (Table S4). Thus, the fraction of human genetic elements that were functionally conserved in chicken was 2.5 times higher for TSSs (21.5%) than for replication origins (8%).

Even among mammals, the conservation of origins is quite limited: 73% of human origins overlapped with human-mouse CGSs (60% expected by chance), but 50% at most were active in mouse (Fig. 6-C). Thus, only 36% of human origins at most appeared to be functionally conserved in mouse.

Finally, for the subset of human origins that were functionally conserved, we examined whether the SNS peak (i.e., the region under selective pressure in humans) corresponded to a zone of high SNS sequence read accumulation in mouse or chicken. As expected, we observed a read enrichment at the homologous position of the SNS peak (Fig. 7). However, the strength of this peak was moderate, which indicated that even for origins with conserved activity, the precise position of the initiation starting point was not conserved. These observations were reproducible over all pairwise comparisons (Fig. S7). Overall, these observations indicate that despite evidence of selective pressure, the location of replication origins is poorly conserved, even on a small evolutionary scale (primate/rodent divergence).

**FIG. 7:**
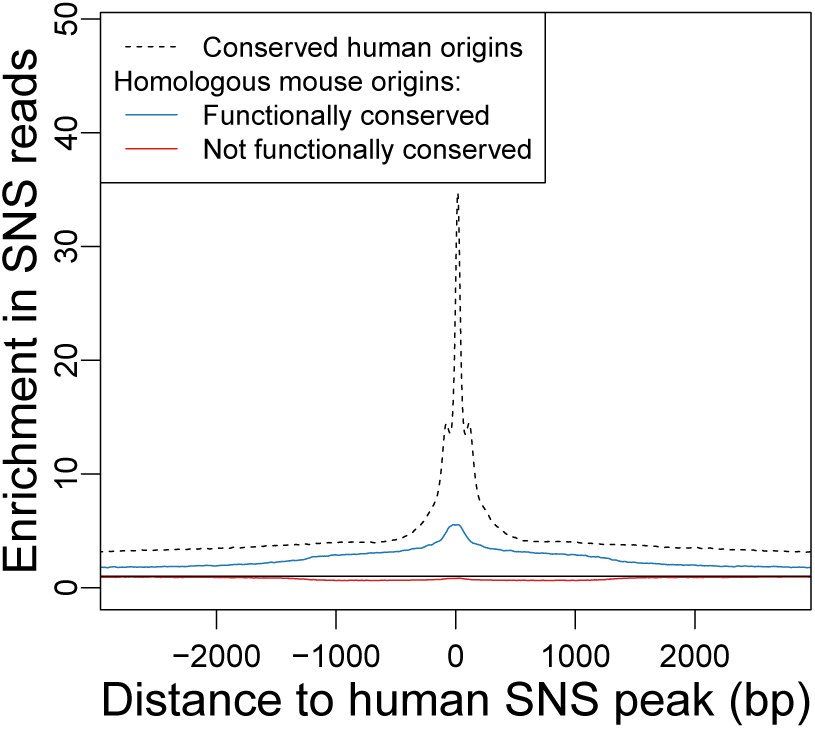
Conservation of replication origin activity at homologous loci. Average SNS read enrichment profile regions centred on the homologue of human SNS peaks in the mouse genome (plain line) compared to the genome-wide average, for conserved origins (*N* = 32, 042) and non-conserved origins (*N* = 56, 078). Dotted line: Average enrichment profile for all human origins overlapping a CGS, centred on the SNS peaks. Only origins with conserved SNS peaks are presented. Black plain line: Expected value without enrichment.

### Long-term conservation of replication origins is driven by associations with functional elements

Mammalian replication origins are known to be frequently associated with TSSs and CGIs (Besnard et al. 2012, Cadoret et al. 2008, Cayrou et al. 2015, Picard et al. 2014, Sequeira-Mendes et al. 2009), and we found the same pattern in chicken (Table S3). Interestingly, the fraction of mammalian origins that were functionally conserved in chicken was 3 to 8 times higher among origins associated with CGI and/or TSS compared to the other origins (Table 2 and Supp. Tab. S5). However, the high conservation level of this subset of origins might simply reflect the selective pressure for promoter function. Alternatively, it is possible that there are specific functional constraints on the subset of promoters that possess replication origin activity. To test this latter hypothesis, we compared profiles of sequence conservation around human TSSs associated or not with an origin. We built these conservation profiles using PhastCons scores based on the multiple alignment of 46 vertebrate species (Siepel et al. 2005). As expected, we found a strong and narrow peak of conservation immediately upstream of the TSS, both for promoters that overlapped a CGI and those that did not (Fig. 8-(A-B)). The average PhastCons scores tended to be slightly higher for promoters with origins than those without but the difference between the two scores was very weak compared to the 4-fold increase in PhastCons scores caused by the presence of TSSs. Moreover, the PhastCons score increase in TSSs associated with origins was independent of the presence of SNS peaks. Indeed, the density of SNS peaks varies substantially along CGI-containing promoters whereas the difference in PhastCons scores between promoters with or without origins remained constant (Fig. 8-A). Thus, the presence of an origin did not impact the level of conservation of promoters.

**TABLE 2:** Conservation of human origins according to their association with TSSs and CGIs: Percentage of origins overlapping conserved genomic segments, and percentage of functionally conserved origins.

|  | Human - Chicken |  | Human - Mouse |  |
| --- | --- | --- | --- | --- |
| Origin Category | % overlapping a CGS | % functionally conserved | overlapping a CGS | % functionally conserved |
| All | 24.8% ($\times 2.1$) | 7.9% ($\times 4.5$) | 72.6% ($\times 1.2$) | 20.7% ($\times 2.4$) |
| CGI[-] TSS[-] | 14.6% | 3.4% | 67.3% | 12.78% |
| CGI[+] TSS[-] | 48.8% | 20.9% | 79.1% | 30.1% |
| CGI[-] TSS[+] | 48.3% | 13.4% | 88.8% | 29.3% |
| CGI[+] TSS[+] | 63.5% | 26.9% | 94.3% | 51.6% |

**FIG. 8:**
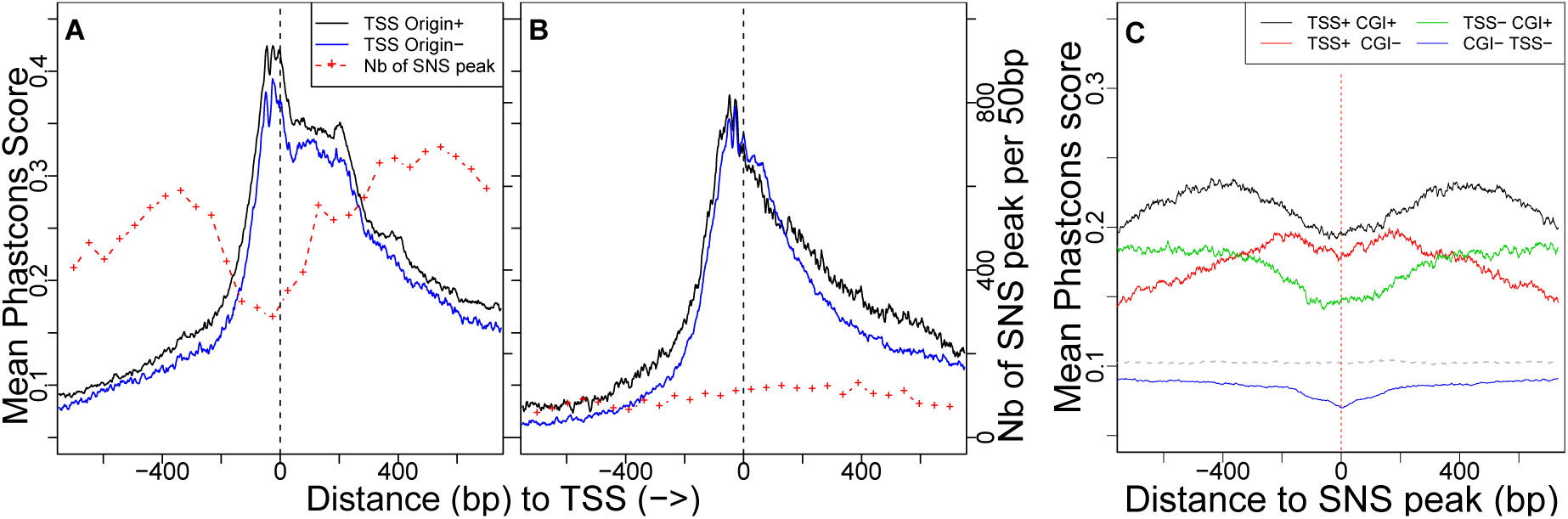
Profile of sequence conservation around TSSs and replication origins. **(A-B)** Human TSSs. Black: TSSs less than 750 bp away from an origin. Blue: TSSs that are not associated with an origin. The red dotted line presents the number of SNS peaks found around all TSSs (smoothed over 50 bp windows). All TSSs are oriented 5′→3′ relative to the transcription unit. **(A)** TSS associated with a CGI **(B)** TSS not associated to a CGI. **(C)** Average PhastCons scores calculated in a 1400 bp region centred on human SNS Peaks, depending on the origin association with TSSs and CGIs. The dashed line presents the genome-wide phastCons score average.

We also examined sequence conservation profiles, centred on SNS peaks of human origins (Fig. 8-C). As expected, replication origins that collocated with TSSs or with CGIs had higher PhastCons scores than those that did not. However, in all cases, the SNS peak position was, on average, less conserved than its immediate surroundings. The same patterns were observed in mouse (Fig. S8). Thus, all these observations indicated that, on the evolutionary scale considered herein, the conservation of replication origins solely resulted from their association with functional elements (CGIs, TSSs) that are under strong selective pressure to be conserved.

## DISCUSSION

Despite the central role of DNA replication in the life cycle of living organisms, the molecular mechanisms inducing the firing of replication are still poorly understood. To track evolutionary signatures that could unravel sequence features controlling replication initiation, we conducted the first comparative analysis of high-resolution maps of replication origins in chicken, human and mouse.

Our meta-analysis first showed that large-scale (100 kb) replication landscapes have been largely remodelled during the evolution of vertebrates. As expected, given the differences in genome sizes, the number of origins is much higher in mammals than in chicken. However, replication landscapes are not simply driven by genome size. In particular, the mouse genome is characterized by a high density of origins of relatively low intensity compared to the two other species (Fig. 2). Thus, whereas the replication initiation activity in human and chicken genomes is concentrated in clusters of very active loci, the mouse genome presents a more uniform distribution.

The analysis of intra-species genetic polymorphism within origins revealed a clear signature of purifying selection, specifically in a narrow region (~40 bp) surrounding the SNS peak, which probably corresponds to the replication initiation site. In the three species, there is a strong excess of origins associated with TSSs or CGIs, which suggests that these genetic elements constitute a favourable context for initiating replication. However, the selection signal on SNS peaks is not driven by functional constraints on TSSs or CGIs since it is observed irrespective of the presence of such elements (Fig. 4). This selective pressure therefore probably results from the presence of specific signals involved in the initiation of replication. Our motif detection procedure revealed an enrichment of CG dinucleotides, CCC triplets and GAG in the 40-bp region surrounding the SNS peaks. Interestingly, this enrichment was observed in all three species. However, these motifs are very short, and hence are also abundant outside of replication origins. This core region must therefore contain another type of signal that does not correspond to simple sequence motifs – for instance, a structural signal. The nature of this signal and its functional role in replication initiation remain to be described. Our analysis also highlighted the heterogeneities of sequence features characterizing this very specific region, as previously shown in mouse origins (Cayrou et al. 2015), hence indicating the potential existence of different classes of origins driven by distinct combination of DNA binding factors (Aladjem and Redon 2017, Cayrou et al. 2015). However, the causes of the observed selective pressure remain enigmatic and might reflect constraints on the local structure of the DNA, rather than on sequence motifs (Comoglio et al. 2015). Intriguingly, we showed that this selective pressure was *not* due to the presence of G4s, which tended to be located further upstream of the SNS peak (Fig. 3). There is evidence that G4s are functionally important, but their presence is not sufficient to initiate replication (Valton et al. 2014).

One striking result of the present analysis is that despite this evidence of purifying selection, which was clearly visible in intra-species polymorphism datasets, inter-species comparisons revealed very limited conservation of replication origins on larger evolutionary scales. Only 4-8% of mammalian origins were functionally conserved in chicken, and 20% of human origins were functionally conserved in mouse. The number of functionally conserved origins ranged from two to four times higher than expected by chance, but this excess could be explained, in large part, by the strong association of origins with other functional elements (TSSs or CGIs) that are themselves conserved (Table 2 and Supp. Tab. S5) in vertebrates. Furthermore, among the small subset of origins that were functionally conserved, the precise location of SNS peaks was not conserved (Fig. 7 and Supp. Fig. S7). All these observations support a rapid turnover of replication start sites during evolution.

One possible hypothesis to reconcile the contrasting patterns of conservation observed on different time-scales is a selective pressure to maintain a sufficient density of replication origins along chromosomes, but no constraint on the precise location of these origins. According to this model, if the number and intensity of origins in a given chromosome are just sufficient to ensure its proper replication, any mutation that would disrupt an origin would be counter-selected. However, a mutation that creates a new origin might occasionally occur. Such a mutation would be neutral and hence could be fixed by random drift. Then, once one additional origin is fixed, mutations that would disrupt a previously existing origin would be neutral. Thus, this model would explain why replication origins appear to be under purifying selection on a short time-scale but not conserved over the long term. A similar model has recently been proposed to explain the evolution of yeast replication origins (Agier et al. 2017). This peculiar temporal conservation pattern is not specific to replication origins: for many functional elements located in non-coding regions, signatures of selection are better revealed by analysis of polymorphisms than by inter-species comparisons (di Iulio et al. 2018).

The high plasticity of the spatial programme of replication is also surprising in light of the evidence showing that the temporal replication programme is organized into large 1 Mb-long domains, which are well conserved in mammals (Ryba et al. 2010, Yaffe et al. 2010).

This discrepancy could imply that the temporal and spatial programme of replication evolve independently. Alternatively, the observation that replication origin density is high in early replication domains and low in late replicating ones could lead to another explanation. Indeed, if origin losses are replaced by origin gains in close proximity, as suggested in a recent study in yeast (Agier et al. 2017), the origin density of replication domains – and hence, their replication timing — would be maintained despite the rapid turnover of origins.

In conclusion, our study highlights the flexibility of the spatial programme of replication in vertebrate genomes, unravelling the existence of a novel genetic determinant of replication origins in vertebrates, the precise nature of which remains to be determined.

## MATERIALS AND METHODS

### SNS data generation

Short nascent strands were purified as previously described (Prioleau et al. 2003). We pooled fractions 15 to 20 containing single-stranded DNA molecules with a size between 1.5 to 2.5 kb in size. We used 500 U of a custom-made *λ*-exonuclease (ThermoFisher Scientific (50 U.ml^-1^)) for each preparation. For the genome-wide mapping of origins, six SNS preparations were obtained independently from 10^8^ cells each and then pooled. SNS were double-stranded by random priming with the Klenow exo-polymerase (# EP0421, Thermo Fisher Scientific) and random primers (#48190011, ThermoFisher Scientific). Adjacent strands were then ligated with Taq DNA Ligase (M0208L, Biolabs). RNA primers were hydrolysed for 30 min at 37°C with 2.5 M NaOH. The reaction was then neutralized with 2.5 M acetic acid, and SNS were purified by phenol chloroform extraction and recovered after ethanol precipitation and resuspension in TE buffer. SNS molecules were quantified with the QBit dsDNA HS assay kit (Q32854, Thermo Fisher Scientific) and 225 ng of double-stranded material was used for fragmentation using a Covaris apparatus with a 50-*μ*l microTUBE-50 AFA Fiber screw-cap tune (PN 520166, Covaris) and a M220 Holder XTU (PN 500488, Covaris). We used a sonication programme with 75 W peak incident power, 10% duty factor and 200 cycles per burst running at 4°C for 195 s to obtain molecules with a size of approximately 180 bp. The library was constructed with the NEBNext©Ultra™ II DNA Library Prep Kit for Illumina©(NEB #E7645S) following the manufacturer’s instructions with some minor modifications. For the adaptor ligation, undiluted adaptor and no size selection were used. The library amplification was performed using NEBNext ©Multiplex Oligos for Illumina©(NEB #E7335S) with NEBNext index 4 primer (#E7314A) and the NEBNext Universal PCR Primer (#E6861A) with three PCR cycles. Library purification was performed with the SPRIselect Reagent kit (Beckman coulter #B23317), and the final elution step was reduced to 20 *μ*l of 0.1X TE. The mean size of the library molecules determined on an Agilent Bioanalyser High Sensitivity DNA chip (5067-4626, Agilent technologies) was 550 bp. Sequencing was performed on a NextSeq 500 Illumina sequencer with a High Output 150 cycles flow cell (paired-end reads of 75 bp) according to standard procedures. A total of 246,14 M clusters were generated and provided 492,28 M of reads. All Illumina sequencing runs were performed at the GENOM’IC facility of the Cochin Institute.

### SNS data meta-analysis

For human and mouse data, we reanalysed SNS replication data generated in previous studies. Human H9, IMR90, HeLa and iPS data were from (Besnard et al. 2012) and K562 data from (Picard et al. 2014). The analysis of mouse data was performed on data from (Cayrou et al. 2015) and (Almeida et al. 2018). For the (Cayrou et al. 2015) dataset, we kept only 2 of the 3 replicates of the experiment (rep1 and rep2) due to the apparent low specificity of data collected for replicate 3 (as confirmed by personal communication with the authors).

### Origins detection

We reanalysed all SNS data using the same bioinformatics procedure. Reads were mapped using Bowtie2 (Langmead and Salzberg 2012) on hg19, mm10 and gal5 genomes. We excluded reads with quality scores lower than 40, as well as those mapped to non-mappable regions of each genome. To detect origins, we applied the methodology from (Picard et al. 2014) with a smaller threshold (√500 bp instead of 2 kb in the original method), as we found that this window size better reflected the outcome of the SNS experiment. To study origins at the base pair scale, we defined “SNS peaks” as the position inside each origin for which the read accumulation profile was maximal. We computed this maximum on a smoothed read accumulation profile using a Gaussian kernel transformation with a bandwidth of ȡ 500 bp.

### Databases

CGI annotations were downloaded from the UCSC database (http://genome.ucsc.edu/index.html) and TSSs from the Eukaryotic Promoter Database (EPD: http://epd.vital-it.ch/) (Dreos et al. 2017) for human and mouse, and from the Ensembl website (http://www.ensembl.org/index.html) (Aken et al. 2016) for chicken.

### Detection of G-quadruplexes

G-quadruplex positions were detected on both strands based on the simple motif G_3_N_1*-*7_G_3_N_1*-*7_G_3_N_1*-*7_G_3_, using a dedicated python script (https://github.com/dariober/bioinformatics-cafe/tree/master/fastaRegexFinder).

### Resampling procedure

To correct for the higher sequencing depth of the chicken SNS experiment, we pooled together all human H9 replicates on the one hand, and all mouse mESC replicates, on the other hand, to generate one large human and one large mouse sample. These two datasets showed a similar sequencing depth of 28 reads per kb in human and 24 reads per kb in mouse. We randomly sampled ~ 20% of the chicken SNS data to generate three independent datasets in the chicken genome with a sequencing depth of 28 reads per kb, comparable to the datasets in human and mouse.

To limit the inter-cell line variability, we focused on the human H9 cell line, because it is the closest cell type to mESCs in mouse for which SNS data are available.

### Cell type specificity

To control our analysis of origin cell type specificity (Fig. 6 (D)), we first pooled and subsampled reads of different replicates of each cell type experiment to generate 5 datasets (one for each cell line) with en equal sequencing depth (32 million mapped reads) and detected origins in these datasets. The sequencing depth of these datasets was much lower than the one previously described such that we in the new H9 set, we retrieved 80% of the H9 origins in the previous set only.

### Base-specific SNP density

The base-specific SNP density shown in Fig. 4 and S4 were defined as the number of observed *x* → *y* variants divided by the observed number of bases *x*, where *x* and *y* are either GC or AT nucleotides.

### Clustering genomic segments based on kmer composition

We sampled random genomic segments of 40 bp in each genome. The number of random segments was the same as the number of origins in each species. Then, the 5-mer frequencies were computed for all random segments. To reduce dimensionality before clustering, we performed dimension reduction based on NMF (non-negative matrix factorization), a modified version of PCA adapted to count data (Gaujoux and Seoighe 2010). Then, we used *k*-means to cluster random segments into homogeneous compositional groups using NMF principal components. The numbers of axes and clusters were chosen by rule of thumb. This automatic rule selected 6 axes. We then detected 6 clusters in the random segments for each species (Supp. Tab. S2), which we referred to as background clusters because they represent the background compositional heterogeneities that characterize each genome. For each species, we also computed the 5-mer frequencies in replication origins, which were assigned to background clusters by using the maximum a posteriori rule based on their kmer composition (Supp. Tab. S2).

### Randomization procedures

For each species, we generated 10 sets of random genomic positions in mappable regions, with the same chromosomal repartition and size distribution as the origins.

### Activity conservation of TSS

To compare TSS conservation to origin conservation, we first extended TSSs on their 5’ side to generate a set of regions with the same size distribution as origins. We then proceeded to a similar conservations analysis on these sets and on CGIs as done for the origins. Random TSS and CGI sets were generated according to the randomization procedure detailed above.

### Homologous scans

To generate homologous scans, we identified the homologs of the SNS peaks in origins overlapping a CGS for all pairwise comparisons. We then computed the number of SNS reads in a window of 3 kb centred on the SNS peak and of its counterpart in the sister genome.

## ACKNOWLEDGEMENTS

M. N. P. and F.P. teams are supported by the Association pour la Recherche sur le Cancer (Labellisation PGA120150202272). F.M., M.-N.P. and F.P. are supported by the Agence Nationale de la Recherche (ANR-15-CE12-0004, OriMolMech). JMF-J and MG are supported by the Spanish Ministry of Economy and Competitiveness (BFU2016-78849-P, co-financed by the European Union FEDER funds). This work was performed using the computing facilities of the computing center LBBE/PRABI.

